# Optimizing real-time phase detection in diverse rhythmic biological signals for phase-specific neuromodulation

**DOI:** 10.1101/2024.08.24.609522

**Authors:** Mengzhan Liufu, Zachary M. Leveroni, Sameera Shridhar, Nan Zhou, Jai Y. Yu

## Abstract

Closed-loop, phase-specific neurostimulation is a powerful method to modulate ongoing brain activity for clinical and research applications. Phase-specific stimulation relies on estimating the phase of an ongoing oscillation in real time and issuing a control command at a target phase. Phase detection algorithms based on Fast Fourier transform (FFT) are widely used due to their computational efficiency and robustness. However, it is unclear how algorithm performance depends on the spectral properties of the input signal and how algorithm parameters can be optimized. We used offline simulation to evaluate the performance of three algorithms (endpoint-corrected Hilbert Transform, Hilbert Transform and phase mapping) on three rhythmic biological signals with distinct spectral properties (rodent hippocampal theta potential, human EEG alpha and human essential tremor). First, we found that algorithm performance was more strongly influenced by signal amplitude and frequency variation compared with signal to noise ratio. Second, our simulations showed that the size of the data window for phase estimation was critical for the performance of FFT-based algorithms, where the optimal data window corresponds to the period of the oscillation. We validated this prediction with real time phase detection of hippocampal theta oscillations in freely behaving rats performing spatial navigation. Our findings define the relationship between signal properties and algorithm performance and provide a convenient method for optimizing FFT-based phase detection algorithms.

## Introduction

Closed-loop, phase-specific neurostimulation is an approach to modulate the nervous system contingent on information from oscillating biological signals detected from the body. This technique has been effectively used in clinical settings to manage a range of neurological disorders and in basic research to investigate the contribution of neural oscillations to brain function (Wendt et al., 2022, Widge, 2024, Widge and Miller, 2019). Phase-specific stimulation systems rely on phase detection algorithms that 1) estimate the phase of an ongoing oscillating signal and 2) issue outputs once a specific phase target has been detected. A commonly used method for phase estimation involves Fast Fourier Transform (FFT)-based methods to extract the frequency band of interest, combined with Hilbert transform to estimate phase. This method has the advantage of simple implementation and fast computation (Wodeyar et al., 2021, Rosenblum et al., 2021). FFT-based methods do not require training in contrast to machine learning and state space modelling approaches (McIntosh and Sajda, 2020, Busch et al., 2022, Wodeyar et al., 2021).

Reliable phase detection is essential for real time applications, but the optimization of algorithm performance faces several challenges since performance depends on the interaction between the algorithm and the input signal. First, it remains unclear how spectral properties of the input signals affect performance. Prior works suggest signal amplitude and signal-to-noise ratio (SNR) are major determinants of algorithm performance (Kim et al., 2023, Shirinpour et al., 2020, Mansouri et al., 2017). However, other work points to temporal properties, such as signal frequency and amplitude variability, being correlated with performance (Mansouri et al., 2017). Thus, the relative contribution of these different signal properties to algorithm performance remains unclear. Second, it is unclear how algorithm performance can be optimized by adjusting algorithm parameters. While signal properties affect the choice of parameters for optimal algorithm performance, the relationship between signal property and optimal algorithm parameter is non-trivial. Identifying the optimal parameters required searching through a large number of possible combinations (Chen et.al 2013).

Our study addresses these challenges. We evaluate the performance of phase detection algorithms in two use cases: generating outputs at a desired target phase consistently over time or at random phases on each cycle. We evaluate the algorithm output with four metrics that quantify how close the algorithm outputs match the desired phase target and the rhythmicity of the output sequences in time. These algorithms are applied to three rhythmic biological signals with diverse spectral properties to determine the relationship between signal properties and algorithm performance. Lastly, we identify optimal algorithm parameters and validate our predictions with a real-time *in vivo* experiment.

## Methods

### Data

Data for our algorithm simulation consists of published datasets (human EEG and essential tremor) and data collected in our lab (rodent LFP). Rodent LFP data was recorded from pyramidal layer of the dorsal hippocampus CA1 region of rats (N=11, male, 44 sessions, 10 minutes each) performing spatial navigation tasks. The data was recorded at a sampling rate of 1500Hz. Human electroencephalogram data (N=40, 40 sessions, 50 minutes each) was recorded at 150Hz from the POZ electrode over the posterior brain of human subjects performing a working memory task (Unsworth et al., 2014). Essential tremor kinematics data (N=11, 346 sessions, 1 minute each) was recorded at 250Hz from an accelerometer attached to the finger of tremor patients receiving non-invasive tremor suppression therapy (Schreglmann et al., 2021).

### Rodent LFP data collection

All procedures were performed under approval by the university’s Institutional Animal Care and Use Committee, according to the guidelines of the Association for Assessment and Accreditation of Laboratory Animal Care. Long Evans rats (N=11, Charles River Laboratories) had adjustable tetrodes targeting the dorsal CA1 region of the hippocampus (AP -3.8, ML -2.75 relative to Bregma). Tetrodes were adjusted into the dorsal CA1 region over two to three weeks after surgical implantation. Animals navigated elevated mazes (Y or F shaped) for reward for 10 minutes (up to 4 session per day, up to 5 days). Data was recorded at 1500Hz using a 128-channel digitizing headstage and the Trodes data recording interface (SpikeGadgets LLC).

### Analysis packages

Analyses were performed in Python based on publicly available libraries. For signal processing, we used Scipy. For statistical analyses we used Numpy, Scipy and Scikit-Learn. For data visualization, we used Matplotlib.

### Phase detection algorithm simulation

We evaluated the performance of three phase detection algorithms with offline simulations. To simulate real time operations, we iterate stepwise across the signal to extract data from a window corresponding to the last N samples (input data window). Data from this window is then processed by the phase detection algorithm to produce an instantaneous phase estimate for the last sample in the data window.

The endpoint-corrected Hilbert transform (ecHT) (Schreglmann et al., 2021) reduces the effect of the Gibb’s phenomenon when estimating the endpoint phase of a signal. It performs fast Fourier transform (FFT) on the raw signal in the data window. The negative frequency components of the Fourier spectrum are eliminated, and the positive frequency components are multiplied by two. The spectrum is then filtered with a causal band pass filter for the frequency range of interest. The processed frequency components are then transformed into the analytical signal using inverse Fourier transform (iFFT). The instantaneous phase at the last sample of the analytical signal is used as the phase estimate.

For Hilbert transform, the input data is first bandpass filtered at the desired frequency range using a forward-backward 2^nd^ order Butterworth filter. We then computed the analytical signal of the filtered signal using Hilbert transform and used the instantaneous phase at the last sample as the phase estimate. The phase mapping algorithm performs linear regression on the bandpass filtered signal and detects a change in the sign of the slope to identify a peak or trough, corresponding to a half cycle of the rhythm. This algorithm assumes a linear relationship between phase and the number of samples elapsed. It calculates the change in phase per signal sample using the duration of the half cycle. The ratio is used to extrapolate the instantaneous phase for future samples, until the next half cycle is reached. We chose a specific target phase and issued an output when the estimated instantaneous phase reached the target phase. To avoid repeated outputs within the same cycle, output was then disabled until the estimated start of the next cycle.

### Algorithm output evaluation

#### Accuracy

Accuracy is defined as the proportion of output events with phases within a π/2 window centered around the target phase.

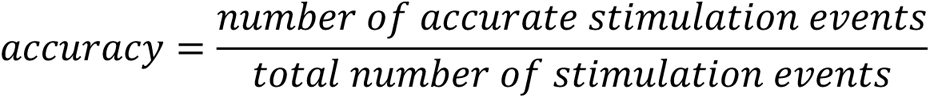

#### Precision

Precision is the resultant vector length of all detected phases. The resultant vector is computed as follows:

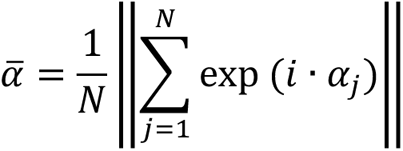

Where N is the number of output events, *α_j_* are the individual detected phases and *α̅* is the resultant phase vector. The resultant vector length is the magnitude of the resultant phase vector *α̅*.

#### Inter-event interval variance

The inter-event interval (IEI) between two consecutive output events is defined as the phase distance between the two events. Variance of the IEI distribution was then calculated as follows:

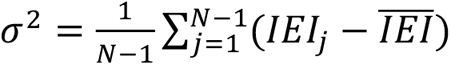

Where N is the number of output events, *IEI_j_* is the inter-event interval between the j-th and the j+1-th event and *IEI̅* is the mean IEI of the output sequence.

#### Autocorrelation

We first compute the autocorrelation for an output sequence within an 80 radians window with 0.2 radian step. We calculated the average autocorrelation values at integer multiples of 2π.

#### Signal property quantification

We measured and compared the following spectral properties of each signal type: signal-to-noise ratio (SNR), amplitude variability and frequency variability. We calculated SNR of each signal by dividing the total power in the frequency band of interest by the total power in all other frequencies. We measured signal amplitude variability using normalized pairwise variability index (nPVI). We extracted the instantaneous amplitude envelope of the target frequency band in the signal. We calculated the mean amplitude for each cycle and calculated the average amplitude difference between consecutive cycles.

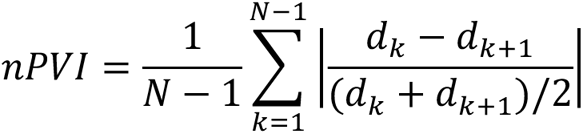

Where N is the total number of cycles in the oscillation, and dk is the average amplitude in cycle k. To quantify the variability in frequency of the signal across time, we calculated the variance of the instantaneous frequency across time. We filtered the signal into the frequency range of interest and used Hilbert transform to extract the instantaneous frequency. The variance of instantaneous frequency across time was then calculated and normalized by the width of the frequency band of interest.

### Simulating the effects of input data window size with sine waves

We quantified the phase estimation error for different input data window sizes with an ideal synthetic sine wave. The sine waves were created at the central frequencies of rodent LFP (8Hz, 1500 samples/s), human EEG (10Hz, 500 samples/s) and essential tremor (8Hz 250 samples/s). Each signal was 20 cycles in length. We used each algorithm to perform phase estimation for each input window size and calculated the circular difference to the actual phase. The estimation error was averaged across all phase estimates.

### Data window size versus algorithm performance simulation

We varied the input data window sizes ranging from 5ms to 500ms in steps of 5ms. For each data input window size, we ran simulations on the actual data using all algorithms to detect the peak of the cycle as the target. We then calculated the accuracy, precision, IEI variance and autocorrelation for each simulation output. This was done using data from 10 sessions from each signal type.

### Random phase generating algorithm simulation

We also simulated algorithms that produce output events at random phases, which we refer to as random phase generating algorithms. The random target algorithm uses the ecHT algorithm to estimate instantaneous phase but selects the output phase target on each cycle by randomly selecting a value from a uniform von Mises distribution (μ=π, κ=0). The random delay algorithm is also based on ecHT but instead produces an output after a random time delay, between 0 and the period of the oscillation, relative to the start of each cycle. The period of the cycle is determined based on the central frequency for the oscillation of interest. The random schedule algorithm involves predefining the timing of the output events relative to the start of the experiment based on the estimated experiment duration and the desired number of output events. This algorithm is blind to the ongoing oscillation and does not perform phase estimation.

### Relating signal property variations to algorithm performance variations

To investigate how signal properties, SNR, and amplitude and frequency variability, relate to algorithm performance, we performed linear regression between each signal property and each algorithm performance metric (Scipy). To understand the relative contributions of these signal properties to algorithm performance variations, we built Generalized Linear Models to model performance using SNR, amplitude variability and frequency variability (Scikit-Learn).

We built the model with data combined across all signal types. All input features to the GLM were z-transformed. We compared the beta coefficients and the feature importance scores of the signal properties.

### *In vivo* testing of optimal window sizes

Two Long Evans rats (male, 45 and 59 weeks old) were implanted with tetrodes targeting the dorsal CA1 region of the hippocampus. We collected data as the animals navigated a Y-shaped maze for reward (15 sessions, 10 minutes each). Data was streamed at 1500Hz using a 128-channel digitizing headstage and the Trodes data recording interface (SpikeGadgets LLC). For real-time signal processing, LFP data from a single channel was streamed to our Python-based ecHT processor. The function detects the peak of the theta oscillation. We concurrently ran four separate, independent instances of the phase detection function, with each instance having a different input data window size. This ensured the same signal was used to test the effects of varying data input window size in real time.

## Results

### Metrics for evaluating algorithm output

When phase detection algorithms are used for phase-specific stimulation, stimulation triggers are issued at a specific phase within each cycle of the oscillation. The goal of a phase-specific stimulation system is to produce sequence of output events that reliably match to the desired phase target for underlying oscillation. The system should consistently produce an output in each cycle, avoiding skipped cycles or repeated outputs within the same cycle (Fig. 1A). The resulting output sequence should form a rhythmic sequence in time that is coupled with the underlying oscillation at the desired target phase.

**Figure 1.**
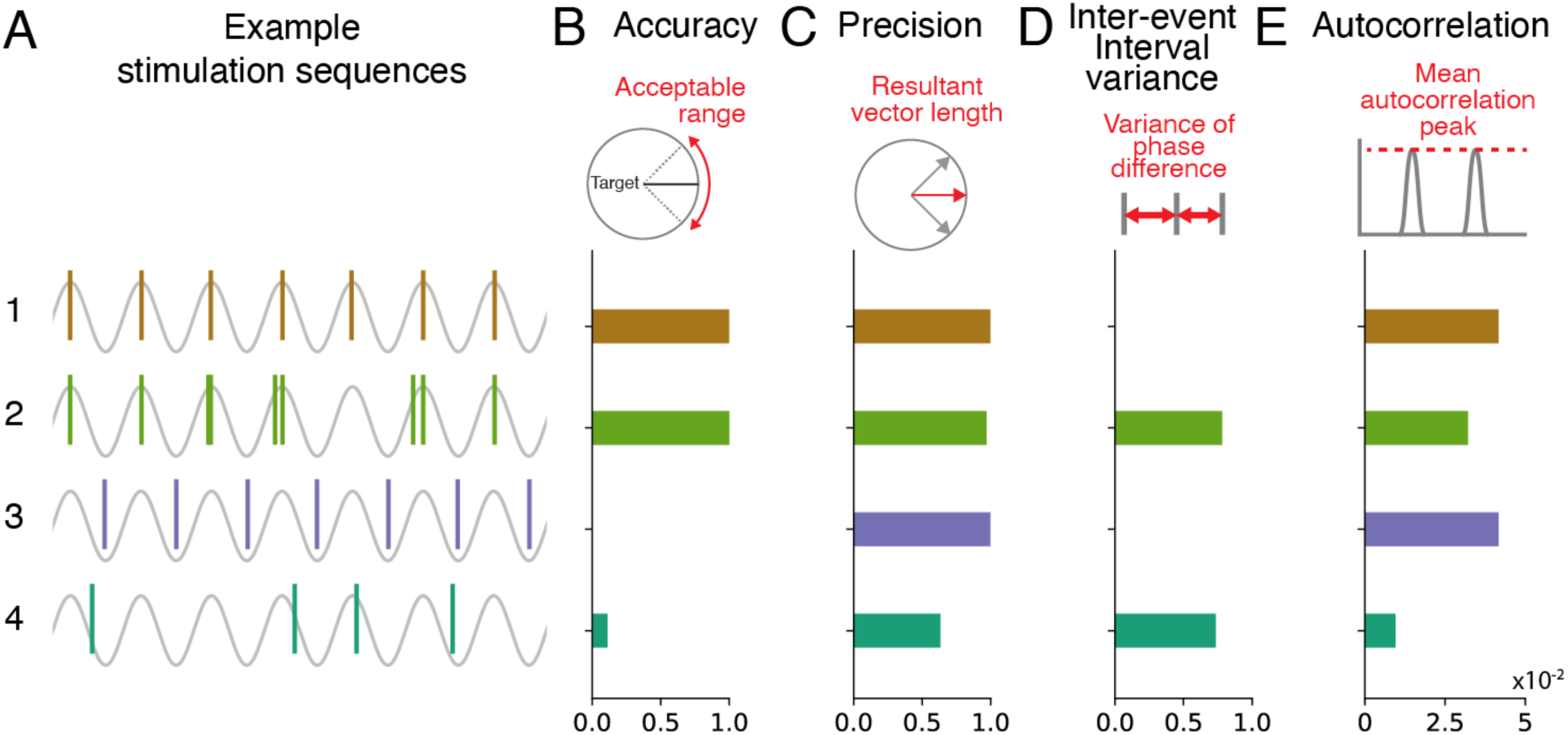
Metrics for evaluating the performance of phase detection algorithms. (A) Example event sequences. 1: consistent events at peak. 2: 20% of the cycles were skipped, repeated events in 30% of the cycles. All events are within a π/2 range centered around peak. 3: consistent events at trough. 4: random events generated from von Mises distributions (μ=π, κ = 0). 10% of the cycles were skipped. (B) Stimulation accuracy quantified by the proportion of events within a π/2 window centered around the target phase (peak). (C) Stimulation precision quantified by the resultant vector length of event phases. (D) Inter-event interval (IEI) variance quantified by the variance of phase differences between events. (E) Autocorrelation quantified by the mean autocorrelation values at integer multiples of 2π over an 80 radians window.

Given the goals, we quantify the performance of the algorithm using four metrics that describe different properties of the output sequence. Two widely used metrics are accuracy and precision. Accuracy reports the proportion of output events within an acceptable range relative to the selected target phase (Fig. 1B). Precision reports the consistency of output events and is quantified using the resultant vector length (Fig. 1C). In addition to quantifying whether outputs are issued at the correct target phase, we chose two metrics to quantify the rhythmicity of the output sequence in time: inter-event interval (IEI) variance and autocorrelation. The inter-event interval (IEI) is the phase difference between two consecutive output events (Fig. 1D). In an ideal case, where each event is delivered consistently at the phase target, the output sequence will have IEIs of 2ρχ. We calculate the variance of the IEIs as a measure of consistency in time. This metric will detect deviations such as multiple output events per cycle or missed events over consecutive cycles. To quantify the rhythmicity of output events, we calculated the mean autocorrelation values of the output event at integer multiples of 2π. Ideal output sequences will have peaks corresponding to multiples of the period (Fig. 1E).

We provide examples to illustrate how these different metrics are necessary to evaluate the algorithm output sequences that could independently vary in phase consistency or rhythmicity. Example sequence 1 is an ideal output for detecting the peak of an oscillation. All the output events are perfectly aligned to the peak, which leads to maximum accuracy, precision and autocorrelation, and no IEI variance. Example sequence 2 has extra, undesired events close to the target phase in some cycles and has missing events in other cycles. In this case, the phase accuracy and precision remain similarly to sequence 1 but IEI variance and autocorrelation reveals decreased rhythmicity. Example sequence 3 is perfectly consistent and rhythmic in time, but all the output events occur at the trough of the cycle which is not the target. This sequence scores high in the rhythmicity metrics but low in accuracy. Example sequence 4 is drawn from a random distribution, the sequence is neither consistent in phase nor rhythmic.

### Evaluating algorithm performance on rhythmic biological signals

We used these metrics to evaluate the performance of three phase detection algorithms: Hilbert transform (Knudsen and Wallis, 2020, Shirinpour et al., 2020, Blackwood et al., 2018, Zrenner et al., 2020, Chen et al., 2013); endpoint-corrected Hilbert transform (Schreglmann et al., 2021, Bressler et al., 2024) (Fig. 2A) and Phase mapping (Fig. 2B). To understand how these algorithms perform on different rhythmic biological signals and how performance depends on signal properties, we selected three biological signals with diverse spectral properties: rodent local field potential (LFP) from the dorsal CA1 region of the hippocampus (Fig. 3A), human electroencephalogram (EEG) from the POZ electrode over the visual regions in posterior brain (Unsworth et al., 2014) (Fig. 3B) and hand kinematics of essential tremor patients measured from an accelerometer attached to the middle finger of their dominant hand (Schreglmann et al., 2021, Brittain et al., 2013) (Fig. 3C). For each signal, we extracted the oscillations in the frequency band with known physiological correlates, tremor frequency for hand kinematics (3-12 Hz) (Schreglmann et al., 2021, Brittain et al., 2013, Cagnan et al., 2013, Welton et al., 2021, Louis and Faust, 2020), alpha rhythm for human EEG (8-12 Hz) (Kim et al., 2023, Bressler et al., 2024, Klimesch et al., 2003, Pantazatos et al., 2023, Peylo et al., 2021, Halgren et al., 2019) and theta rhythm for rat LFP (6-9 Hz) (Siegle and Wilson, 2014, Hyman et al., 2003, Lurie et al., 2022, Buzsáki, 2002, Buzsáki and Moser, 2013) (Fig. 3D). These oscillations differ in signal to noise ratio (SNR) (Kruskal-Wallis test, H (2, 424) = 196, p < 1e-42), amplitude variability over time as measured by nPVI (Kruskal-Wallis test, H (2, 424) = 15.7, p < 1e-21), and frequency variability (Kruskal-Wallis test, H (2, 424) = 101, p < 1e-3) (Fig. 3E – 3G).

**Figure 2.**
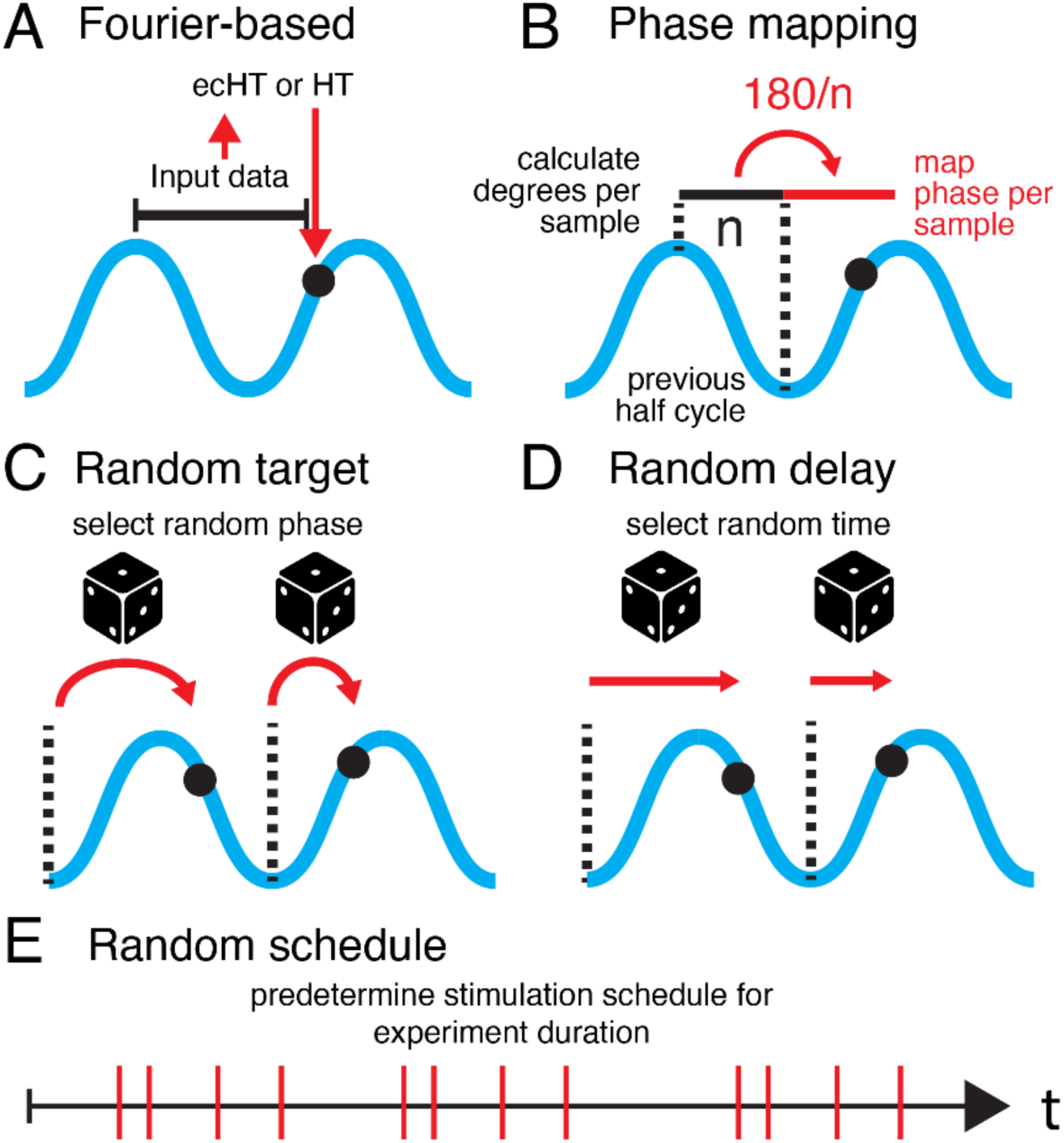
Phase detection and random phase generating algorithms. (A) Schematic of Fourier-based ecHT and HT algorithms. (B) Schematic of the PM algorithm. (C) Schematic of the random target algorithm. (D) Schematic of the random delay algorithm. (E) Schematic of the random schedule algorithm.

**Figure 3.**
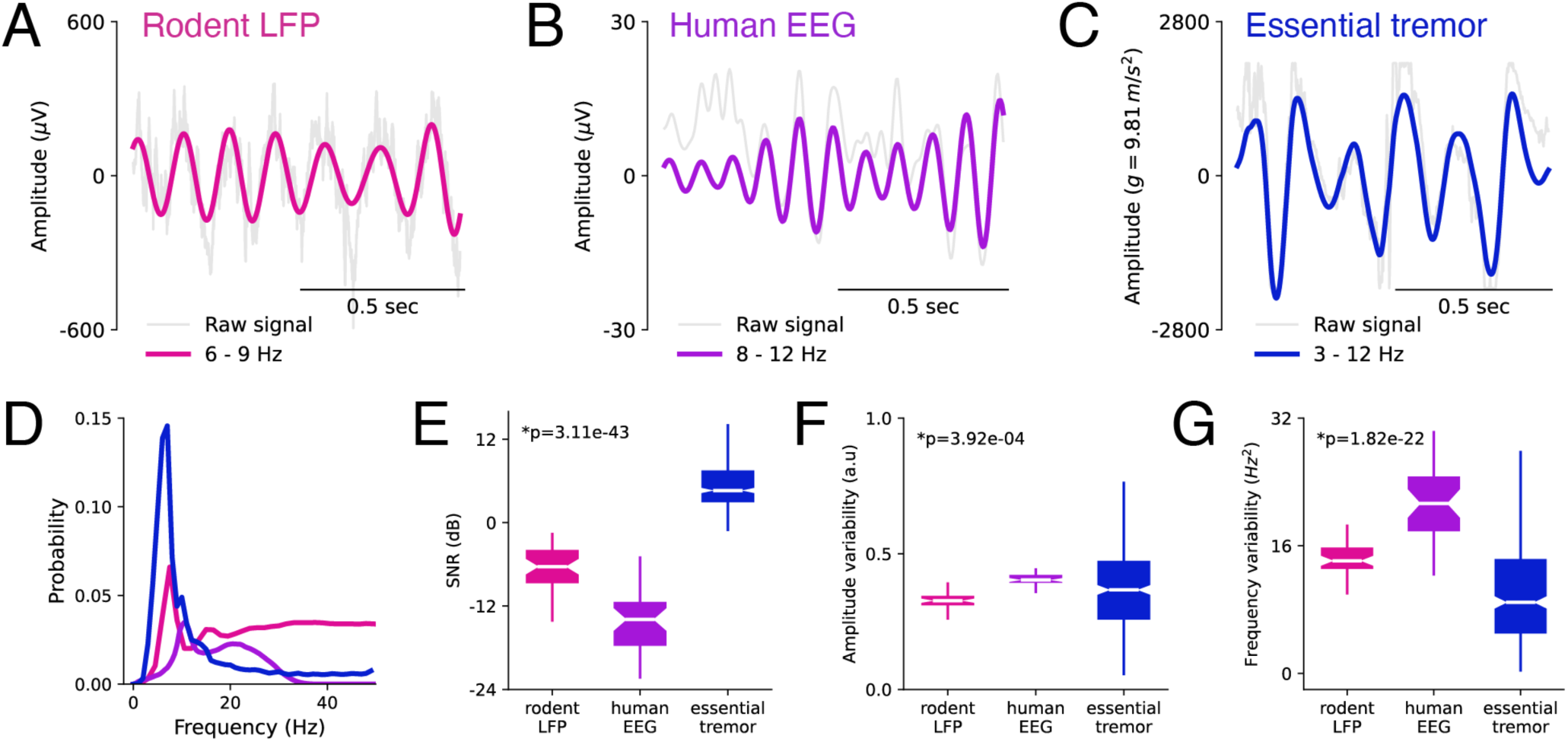
Spectral properties for three rhythmic biological signals. (A) Rodent LFP from the dorsal CA1 of the hippocampus, filtered in the theta range (6-9Hz). (B) Human EEG from the POZ electrode, filtered in the alpha range (8-12Hz). (C) Hand acceleration from human essential tremor patient, filtered in the tremor range (3-12Hz). (D) Spectrograms of sample signals (mean). (E) Box plot of signal-to-noise ratio (SNR) distributions. Statistical difference across signal type was tested using Kruskal-Wallis test (h=1.96e2, p=3.11e-43). (F) Box plot of amplitude variability distributions. Statistical difference across signal type was tested using Kruskal-Wallis test (h=1.57e1, p=3.92e-4). (G) Box plot of frequency variability distributions. Statistical difference across signal type was tested using Kruskal-Wallis test (h=1.00e2, p=1.82e-22).

To accurately compare algorithm performance across signal types, we first optimized the performance of each algorithm on each signal type. Previous work pointed to the input data window size being an important factor in algorithm performance, but it is unclear how the optimal window size can be determined (Shirinpour et al., 2020). The input data window size is the number of data points in the past used for estimating the current phase. We hypothesized that input data window size may affect the performance of the algorithms in two ways. First, window size determines whether data in the window spans complete oscillatory cycles. FFT-based phase estimation is sensitive to input data spanning less than complete cycles (Schreglmann et al., 2021). Using simulations with sine waves, we found that the estimation errors using HT is strongly modulated by window size (Fig. 4A, D and G). In contrast, ecHT is less sensitive to window size. We found that estimation error can be reduced when the data window size is close to multiples of the oscillation period. Phase mapping is the least sensitive to window size since it does not rely on Fourier transform or Hilbert transform.

**Figure 4.**
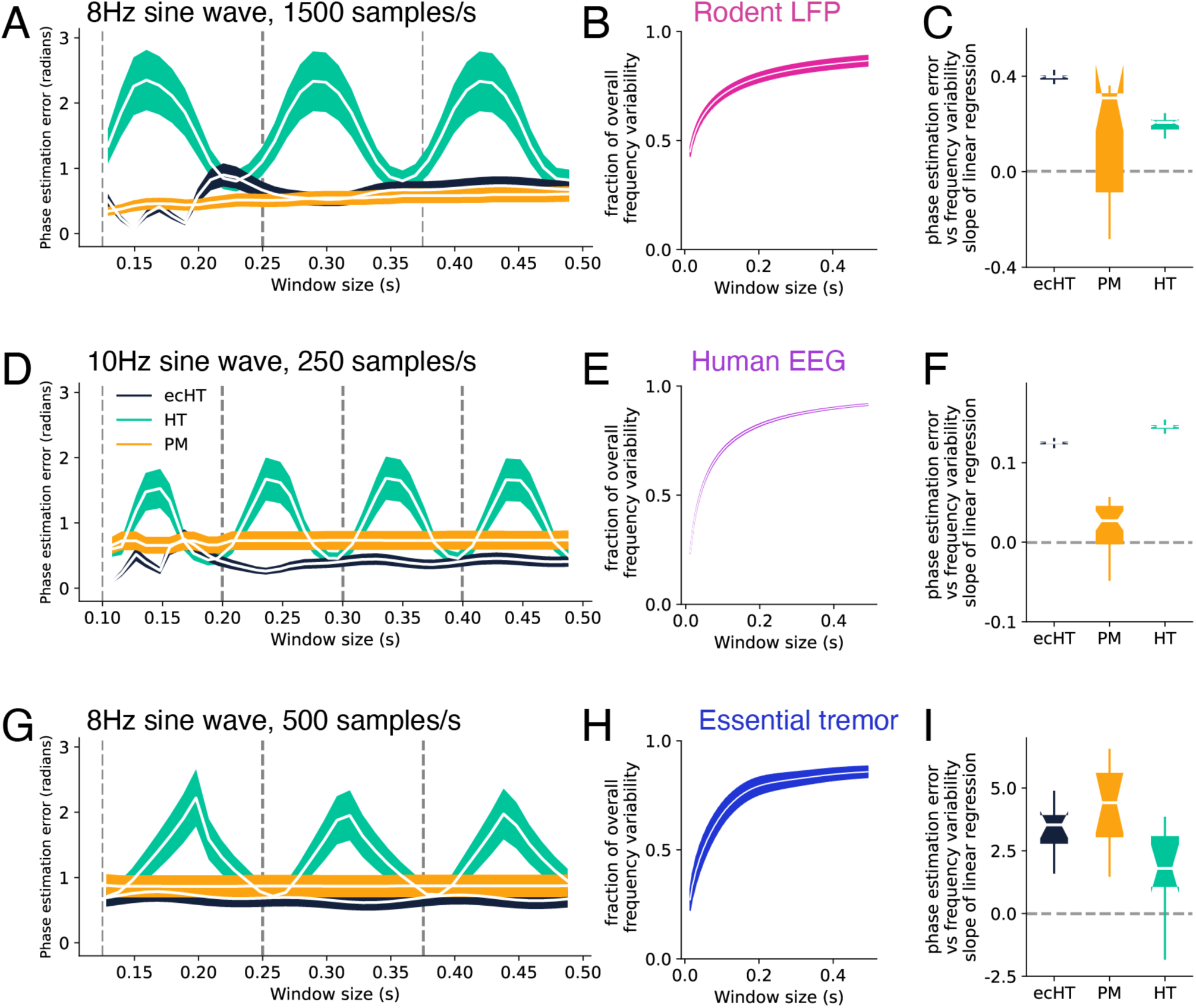
Input data window size and signal frequency variability independently affect algorithm performance. (A) Algorithm phase estimation error (mean±s.e.m.) for 8Hz sine wave at 1500 samples/s to match sampling rate for rodent LFP signal. (B) Rodent LFP signal variability (mean±s.e.m.). (C) Rodent LFP data. Slope of linear regression between estimation error and frequency variability within each data input window (125ms) for each algorithm. Statistical differences between distribution means and zero were tested using Wilcoxon signed-rank test (ecHT: W=253, p=2.38e-07; HT: W=266, p=2.29e-07; PM: W=184, p=3.17e-02). (D) Algorithm phase estimation error (mean±s.e.m.) for 10Hz sine wave at 250 samples/s to match sampling rate for human EEG signal. (E) Human EEG signal variability (mean±s.e.m.). (F) Human EEG data. Slope of linear regression between estimation error and frequency variability within each data input window (100ms) for each algorithm. Statistical difference between distribution means and zero were tested using Wilcoxon signed-rank test (ecHT: W=666, p=1.46e-11; HT: W=721, p=1.83e-08; PM: W=452, p=3.10e-02). (G) Algorithm phase estimation error (mean±s.e.m.) for 8Hz sine wave at 500 samples/s to match sampling rate for essential tremor signal. (H) Essential tremor signal variability (mean±s.e.m.). (I) Essential tremor data. Slope of linear regression between estimation error and frequency variability within each data input window (125ms) for each algorithm. Statistical differences between distribution means and zero were tested using Wilcoxon signed-rank test (ecHT: W=66, p=4.88e-4; HT: W=75, p=7.34e-04; PM: W=452, p=3.10e-02).

In addition, window size also determines the frequency variability of the data in the window (Fig. 4B, E and H). Previous work found that the frequency variability of the signal across an entire session is negatively correlated with algorithm performance (Shirinpour et al., 2020). Statistically, the variability of data within a window will approach the variability of the entire signal as the length of the window increases. Because window size also affects the fraction of full cycles contained within each window, we ask whether these two factors affect algorithm performance independently. We ran simulations for each algorithm on each type of the three biological signals by fixing the data window size at the duration of one cycle. We then correlated the frequency variability with phase estimation error for each iteration across the whole signal. We found consistently positive correlation for all algorithms and signal types (Fig. 4C, F and I). Thus, higher frequency variability found in longer input data windows will lead to higher phase estimation errors.

Given that these two factors that independently affect algorithm performance, we cannot assume the optimal window size estimated from sine wave simulations can be extrapolated to real data. Therefore, we tested algorithm performance on the three biological signals across a range of window sizes and found that optimal window size corresponds approximately to the length of one cycle of the oscillation. For each algorithm, we calculated the performance metrics for simulations with different window sizes ranging from 50ms up to 500ms. We found that the performance metrics of ecHT and HT varied across this range and differed between signal types (Fig. 5A – 5D). As expected, the performance of PM was consistent across window sizes since it does not require time-frequency transformation. The window size yielding the best accuracy, precision and autocorrelation was close to the cycle length (Fig. 5E). The optimal window size for reduced IEI variance was slightly above twice the cycle length. Further, we observed periodicity in the performance metrics, where local maxima occurred at multiples of the cycle length, consistent with our sine wave simulations (Fig. 4). We found differences across signals in how well cycle length correlates with the optimal window size. The window size for optimal algorithm performance is closest to the cycle length for the rodent LFP and human EEG signals, compared with the human essential tremor signals. These results point to a trade-off between phase reliability versus temporal consistency in the algorithm output. Input data windows containing one cycle of signal produce optimal accuracy, precision and autocorrelation, but slightly longer windows minimize IEI variance. Input windows containing more than two cycles of signal did not produce performance benefits.

**Figure 5.**
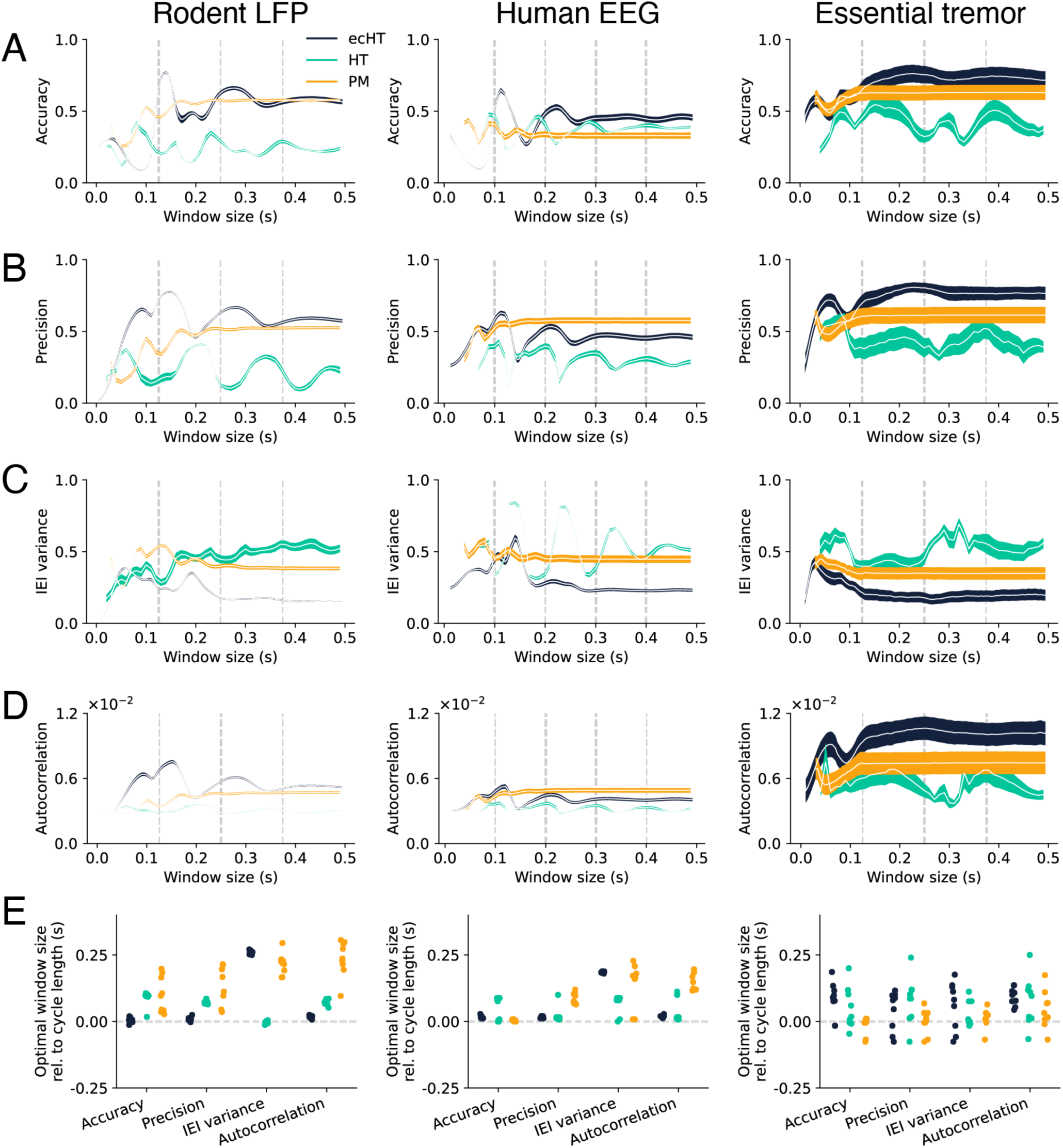
Relationship between algorithm performance and data input window size. (A) Accuracy of algorithm output (ecHT: black, HT: green, and PM: yellow) for rodent LFP (left), human EEG (center) and essential tremor (right) data. Mean and standard error of the mean are shown for each algorithm. (B) Precision of algorithm output. (C) IEI variance of algorithm output. (D) Autocorrelation of algorithm output. (E) Difference between window size with optimal performance metric value and the duration of one oscillation cycle for each algorithm (ecHT: black, HT: green, and PM: yellow) and signal type, rodent LFP (left), human EEG (center) and essential tremor (right). Each point represents data from one individual.

Having determined the optimal window size for each algorithm, we then compared their performance across signals. We ran simulations using each algorithm to detect four evenly distributed phase targets across the cycle, -π/2 (rising), 0 (peak), π/2 (falling), π (trough) (Fig. 6). For each algorithm, performance differed across signal types (Table 1, p-value of signal type < 0.05 for all metrics). Rodent LFP and human EEG data yielded the most consistent outcomes across individual subjects for all algorithms (Levene’s test, F (2, 424) > 10, p < 1e-5 for all metrics. p < 1e-2 for all post-hoc pairwise comparisons). There was an overall performance difference between the algorithms (Table 1, p-value of algorithm < 5e-2 for all metrics), with ecHT yielding better performance compared with PM or HT across all metrics (Tukey’s HSD test, p < 1e-2 for all pairwise comparisons). Algorithm performance varied across the four phase targets, with ecHT being the most consistent across target phases.

**Figure 6.**
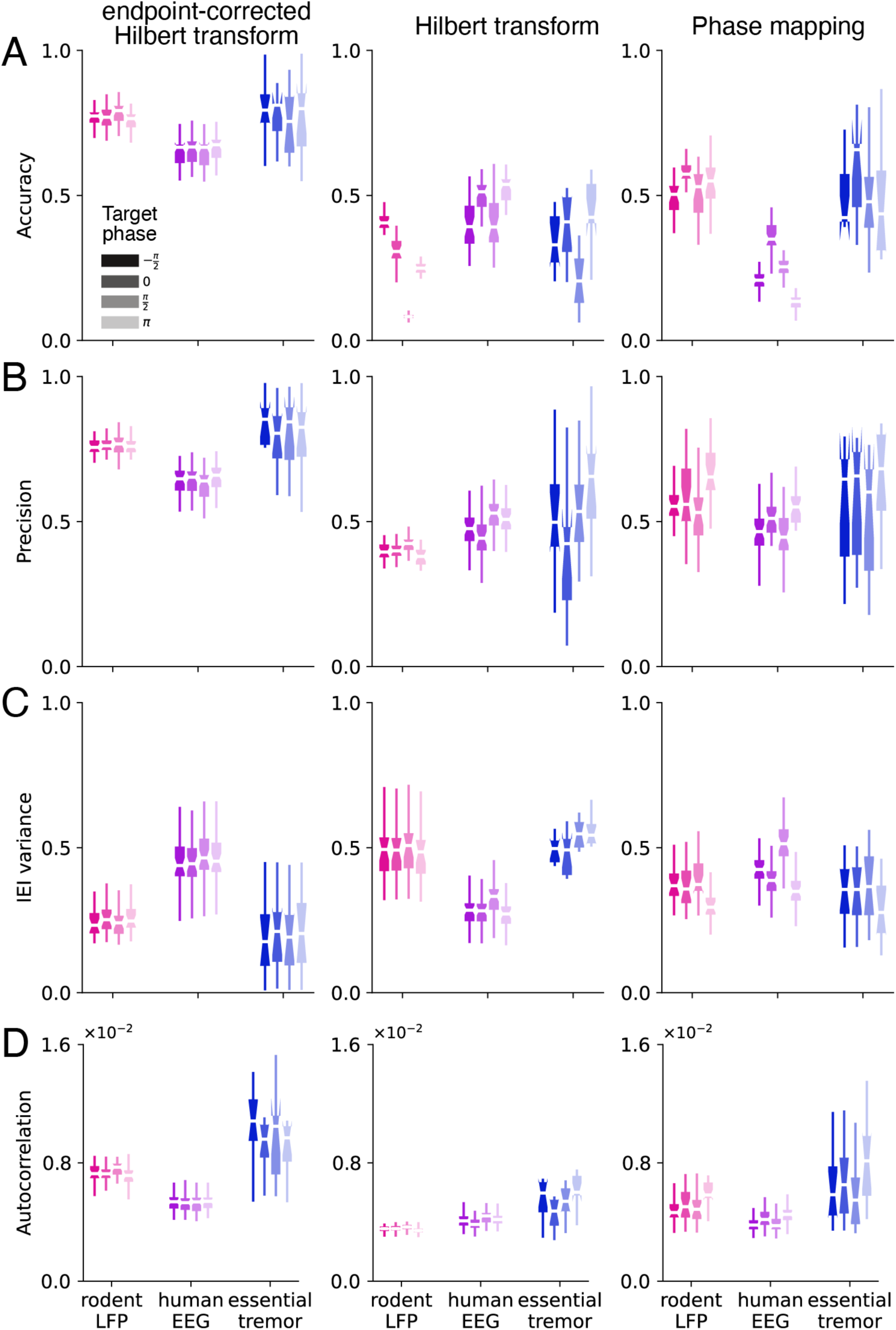
Algorithm performance depends on signal type. (A) Accuracy of algorithm output for each signal type (rodent LFP: pink, human EEG: purple, essential tremor: blue) and algorithm (ecHT: left, HT: center, PM: right). Each algorithm was run for four target phases (-ρχ/2, 0, ρχ/2, ρχ). Shading indicates the target phase for each. (B) Precision of algorithm output. (C) IEI variance of algorithm output. (D) Autocorrelation of algorithm output for each signal type and algorithm when input data window size is fixed at the optimum.

**Table 1.** Three-way ANOVA of the effects of signal type, algorithm and target phase on each algorithm performance metric.

| *Performance metric* | *Source of variation* | *df* | *SS* | *MS* | *F* | *p-value* |
| --- | --- | --- | --- | --- | --- | --- |
| *Accuracy* | Target phase | 3 | 0.7356 | 0.2452 | 30.3902 | 1.33e-18 |
|  | Algorithm | 2 | 19.9466 | 9.9733 | 1236.1771 | 2.70e-244 |
|  | Signal type | 2 | 1.5098 | 0.7549 | 93.5673 | 3.32e-37 |
| *Precision* | Target phase | 3 | 0.2583 | 0.0861 | 9.7180 | 2.65e-06 |
|  | Algorithm | 2 | 8.3663 | 4.1831 | 472.2083 | 9.82e-136 |
|  | Signal type | 2 | 0.8372 | 0.4186 | 47.2504 | 4.11e-20 |
| *IEI variance* | Target phase | 3 | 0.3197 | 0.1066 | 14.6642 | 2.66e-09 |
|  | Algorithm | 2 | 0.2262 | 0.1131 | 15.5641 | 2.34e-07 |
|  | Signal type | 2 | 0.3100 | 0.1550 | 21.3322 | 9.47e-10 |
| *Autocorrelation* | Target phase | 3 | 1.7e-06 | 6.0e-07 | 3.0279 | 2.87e-02 |
|  | Algorithm | 2 | 8.64e-05 | 4.32e-05 | 234.2373 | 1.21e-80 |
|  | Signal type | 2 | 1.02e-04 | 5.11e-05 | 276.8501 | 6.78e-92 |

We next analyzed the performance of algorithms that generate outputs without phase specificity, which we refer to as random phase generating algorithms. Laboratory experiments using phase specific stimulation often require a non-phase specific stimulation condition as control. The control helps establish that the observed effects are specific to a unique stimulation phase for the rhythmic signal of interest (Knudsen and Wallis, 2020, Xie et al., 2023, Ngo et al., 2013, West et al., 2022, Hampson et al., 2018, Roeder et al., 2024, Kanta et al., 2019, Zanos et al., 2018, Dong et al., 2023). We evaluated the output of three random phase generating algorithms (random target, random delay and random schedule) (Fig. 2C – E, Fig. 7). There was an overall difference between their outputs (Table 2, p-value of algorithm < 0.05 for all metrics). Random target produced the most random output phases as measured by resultant vector length (Tukey’s HSD test, p < 0.05 for all pairwise comparisons) (Fig. 7A). Random schedule and random target produced the most arrhythmic sequences as measured by IEI variance and autocorrelation (Tukey’s HSD test, p < 0.05 for all pairwise comparisons) (Fig. 7B, 7C). The algorithm performance was again dependent on signal type (Table 2, p-value of signal type < 0.05 for all metrics) with rodent LFP and human EEG yielding the most random and arhythmic outcomes. Essential tremor signals yielded less random output overall (Tukey’s HSD test, p < 0.05 for all pairwise comparisons) and less consistent autocorrelation across individuals (Levene’s test, F (2, 424) = 21.08, p = 6.73e-09. p < 0.05 for all post-hoc pairwise comparisons).

**Figure 7.**
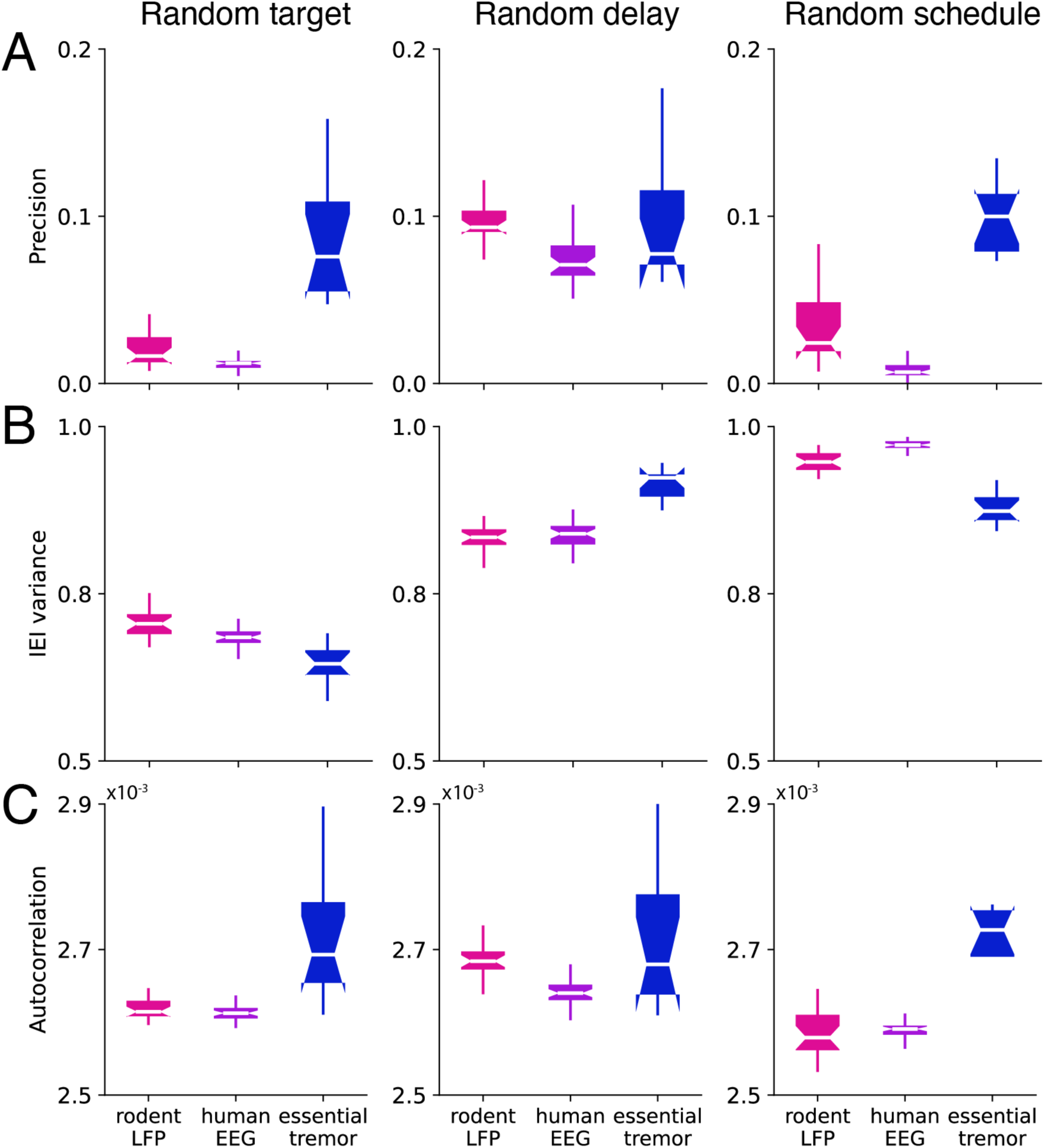
Random phase generating algorithm performance depends on signal type. (A) Precision of algorithm output for each signal type (rodent LFP: pink, human EEG: purple, essential tremor: blue) and random phase generating algorithm (random target: left, random delay: center, random schedule: right). (B) IEI variance of algorithm output. (C) Autocorrelation of algorithm output.

**Table 2.** Two-way ANOVA of the effects of signal type and algorithm on each algorithm performance metric.

| *Performance metric* | *Source of variation* | *df* | *SS* | *MS* | *F* | *p-value* |
| --- | --- | --- | --- | --- | --- | --- |
| *Precision* | Algorithm | 2 | 0.1447 | 0.0724 | 150.4087 | 1.88e-40 |
|  | Signal type | 2 | 0.1124 | 0.0562 | 116.8723 | 3.04e-34 |
| *IEI variance* | Algorithm | 2 | 2.3482 | 1.1741 | 2062.7140 | 2.65e-133 |
|  | Signal type | 2 | 0.0097 | 0.0049 | 8.5439 | 2.76e-04 |
| *Autocorrelation* | Algorithm | 2 | 1.92e-07 | 5.46e-08 | 14.4489 | 1.40e-06 |
|  | Signal type | 2 | 3.16e-07 | 1.58e-07 | 41.7439 | 7.90e-16 |

### Signal properties affect algorithm performance

We next asked how the spectral properties of a signal are correlated with algorithm performance. We performed linear regression between each signal property and each algorithm performance metric (Supp. Fig. 1, 2, 3). For all signal types and phase-specific algorithms, SNR was positively correlated to algorithm performance, and amplitude and frequency variability were negatively correlated to performance. For all signal types, the performance variations of ecHT consistently showed strong correlations with amplitude and frequency variability. For PM and HT, signal properties were less correlated with performance metrics. In contrast to previous findings indicating SNR is a strong determinant of algorithm performance (Kim et al., 2023, Wodeyar et al., 2021, Shirinpour et al., 2020, Zrenner et al., 2020, Rodriguez Rivero and Ditterich, 2021), we found SNR was least correlated with performance across several metrics.

To understand the relative contribution of these signal properties to algorithm performance variations, we built Generalized Linear Models to model algorithm performance across signal types using all three signal properties: SNR, amplitude variability and frequency variability. We examined the beta coefficients and calculated feature importance scores for each of these properties. There was an overall difference in the relative importance of the signal properties across all algorithms (Supplementary Table S1, S3; beta coefficients: p = 6.57e-3 for signal features; importance scores: p = 7e-6 for signal features) (Fig. 8A). Signal amplitude and frequency variability impact algorithm performance more than SNR. Across signal types and algorithms, signal amplitude and frequency variability had higher coefficients and importance scores compared with SNR. Amplitude variability had either comparable or higher feature importance than frequency variability (see Supplementary Table S2, S4). Compared with ecHT and HT, feature importance differences between signal features were less significant for PM (p > 0.05 for all pairwise comparisons). These relationships are evident when comparing the amplitude and frequency variability of each data type (Fig. 3F-G) with their respective ecHT performance (Fig. 6, left column)

**Figure 8.**
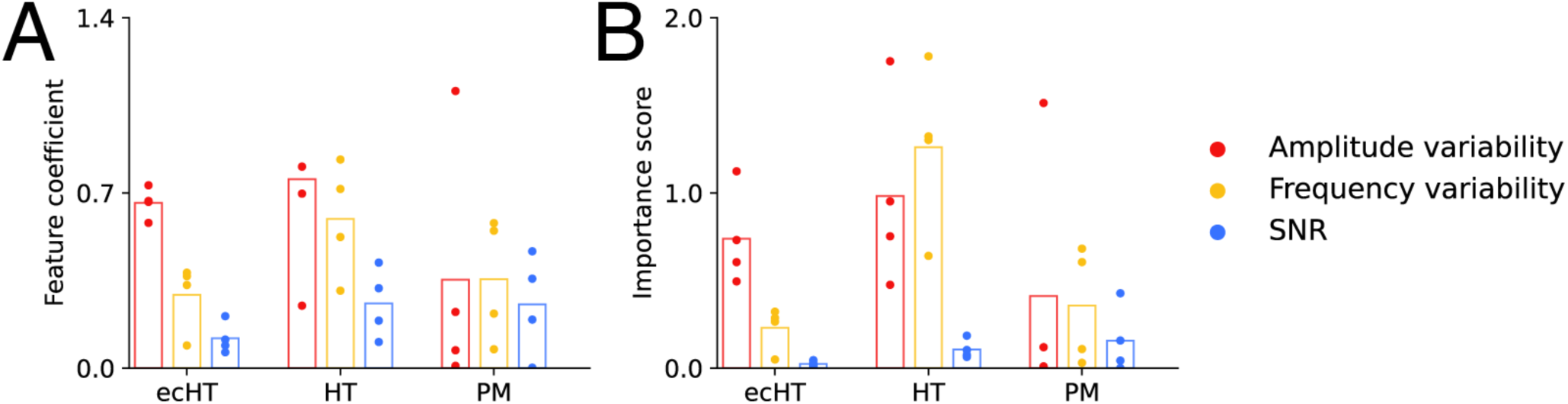
Signal amplitude and frequency variability contribute to algorithm performance. (A) GLM models were constructed with amplitude variability (red), frequency variability (yellow) and SNR (blue) as features to predict algorithm performance across all three signal types. The plot shows the beta coefficients for each input feature. Each data point shows the beta coefficient for one performance metric. The mean beta coefficients are indicated by the vertical bars. Statistical difference across features was tested using two-way ANOVA (p = 6.57e-3 for signal features) and Tukey’s Honest Significant Difference (HSD) test for post-hoc testing (refer to Supplementary Table S2). (B) The plot shows the feature importance scores for each input feature. Each data point shows the importance score for one performance metric. The mean importance scores are indicated by the vertical bars. Statistical difference across features was tested using two-way ANOVA (Importance score: p = 7e-6 for signal features) and Tukey’s Honest Significant Difference (HSD) test for post-hoc testing (refer to Supplementary Table S4).

### *In vivo* validation of algorithm optimization

Our simulation results predict that Fourier-based phase estimation can be optimized by matching the input data window size with the period of the oscillation (Fig. 5). We performed a real time *in vivo* experiment to validate this prediction (Fig. 9A). We selected the ecHT algorithm since it performed consistently better compared with HT or PM. We selected four window sizes, 80ms, 150ms, 200ms and 280ms, corresponding to 0.5, 1, 1.5 and 2 cycles of the theta oscillation respectively. Based on our simulation data (Fig. 5 Rodent LFP), these window sizes should produce distinct performance results. We applied ecHT to detect the peak of the theta oscillation in the dorsal CA1 region of the hippocampus in two rats as they navigated a Y maze for reward.

**Figure 9.**
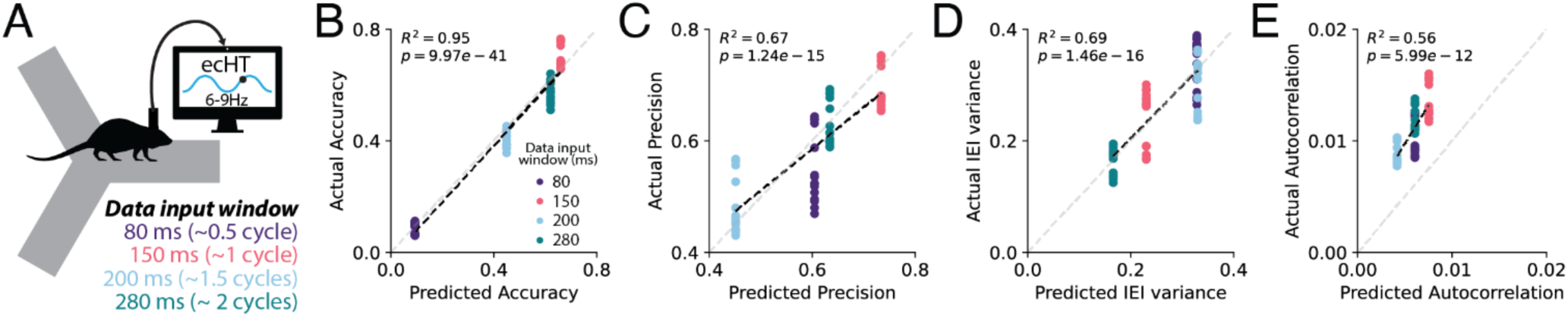
*In vivo* validation of ecHT algorithm performance for four data input window sizes. (A) Schematic of *in vivo* real-time experiment. Two rats were implanted with recording electrodes in the dorsal CA1 region. The ecHT algorithm was used for real-time phase detection in the theta frequency band (6 – 9 Hz) as animals ran on a Y maze. (B) Actual versus predicted accuracy for four different input data window sizes (80ms: purple, 150ms: pink, 200ms: blue, 280ms: green). Each point represents results from one 10 minute session. Linear regression (R^2^=0.95, p=9.97e-41). (C) Actual versus predicted precision for four different input data window sizes. Linear regression (R^2^=0.67, p=1.25e-15). (D) Actual versus predicted interevent interval for four different input data window sizes. Linear regression (R^2^=0.69, p=1.46e-16). (E) Actual versus predicted autocorrelation for four different input data window sizes. Linear regression (R^2^=0.56, p=5.99e-12).

The observed real-time performance was highly correlated with our predictions (accuracy: R^2^ = 0.95, p=9.97e-41; precision: R^2^ = 0.67, p = 1.24e-15; IEI variance: R^2^ = 0.69, p = 1.46e-16; autocorrelation: R^2^ = 0.56, p = 5.99e-12). The best performance across metrics was observed for an input data window corresponding to approximately 1 cycle of the oscillation, followed by 2 cycles. The other window sizes, 0.5 and 1.5 cycles, performed less well. For accuracy, an input data window size to 1.5 cycles reduced the performance by half compared with a window size of 1 cycle (Fig. 9B), pointing to the importance of matching the window size to the period of the oscillation.

## Discussion

Closed-loop phase-specific neurostimulation is a powerful method to modulate ongoing brain activity for clinical applications and basic research (Widge and Miller, 2019, Wendt et al., 2022, Widge, 2024). For clinical applications, phase-specific stimulation has been used to terminate epileptic seizures (Takeuchi et al., 2021), suppress essential tremor (Cagnan et al., 2013, Brittain et al., 2013, Schreglmann et al., 2021), augment sedation (Smith et al., 2024) and modulate sleep patterns (Bressler et al., 2023, Bressler et al., 2024, Dong et al., 2023). In basic neuroscience research, phase-locked stimulation has been used to modulate corticospinal excitability and synaptic plasticity (Zanos et al., 2018, Zrenner et al., 2023, Wischnewski et al., 2022), investigate phase-specific function of neural oscillations in cognition (Ngo et al., 2013, Siegle and Wilson, 2014, Kanta et al., 2019, Mankin and Fried, 2020, Lurie et al., 2022, Xie et al., 2023, Knudsen and Wallis, 2020). Optimizing the performance of online phase detection algorithms and closed-loop phase-specific stimulation systems will benefit these applications.

Our work identifies one method for optimizing the performance of Fourier transform-based algorithms that is convenient to implement. We found performance can be optimized by adjusting the input data window to match the duration of one cycle of the oscillation. This was shown in simulations and validated with an *in vivo* real-time experiment in rats. Our simulation work focuses on theta rhythm in rodent, alpha rhythm and essential tremor in humans, whether our method can generalize to higher frequency oscillations is a topic for future investigation. Higher frequency oscillations such as beta (12-30Hz) and gamma (30-90Hz) oscillation may have different spectral properties and may require further optimization.

Whether different algorithm classes are sensitive to input data window size remains to be determined. We found phase mapping algorithm is less sensitive to window size as it does not involve time-frequency transformation. Phase estimation algorithms using machine learning and state space modelling approaches also do not involve time-frequency transformation, and thus may be less dependent on window size (Wodeyar et al., 2021, Busch et al., 2022, Rosenblum et al., 2021, McIntosh and Sajda, 2020, Tseng et al., 2023). Additional classes of phase estimation algorithms include autoregressive forecast to address the Gibbs phenomenon seen in online filtering (Chen et al., 2013, Blackwood et al., 2018, Schatza et al., 2022, Onojima and Kitajo, 2021), Kalmann filtering to adjust frequency filter range in real time or Genetic algorithms to search for a set of globally optimal parameters (Van Zaen et al., 2010, Chen et al., 2013, Rodriguez Rivero and Ditterich, 2021).

In addition to phase-specific stimulation, many closed-loop laboratory experiments involve a non-phase specific condition as control. The objective can be to ensure that the effects observed with phase specific stimulation are unique to the phase of the stimulation (Lurie et al., 2022, Chen et al., 2024, Knudsen and Wallis, 2020, Cagnan et al., 2017). This objective requires a similar number of stimulations to be delivered but without phase preference. In other studies, the objective is instead to eliminate the rhythmicity of the underlying neural activity by delivering arhythmic stimulation sequences (Kanta et al., 2019, Schreglmann et al., 2021). Non-phase specific stimulation approaches include open-loop stimulation at fixed time intervals (Xie et al., 2023, Ngo et al., 2013, West et al., 2022, Schreglmann et al., 2021), stimulation at randomly determined time points (Hampson et al., 2018, Kanta et al., 2019, Knudsen and Wallis, 2020, Roeder et al., 2024) and replaying phase-specific stimulation sequence at a later time in the same brain region (Zanos et al., 2018, West et al., 2022). Although closed-loop random phase generation approach produces greater phase randomness than the open loop approach, the open loop approach produces more temporally randomized phase output sequences. These results indicate that different random phase generating algorithms vary in the randomness of their output, and the choice of the control algorithm needs to be tailored to the experiment.

We also identified frequency and amplitude variability as two spectral properties of the input signals that impair algorithm performance. For each iteration of phase estimation, we found that the estimation error is correlated with frequency variability within the data input window. Thus, oscillations with high frequency variability may pose a challenge for phase estimation using FFT-based methods. Amplitude and frequency variation negatively impacted algorithm performance, while SNR had a positive impact on algorithm performance. Comparatively, amplitude and frequency variation had stronger influence on algorithm performance than SNR, which has not been previously reported. Thus, optimal performance for an algorithm depends on the properties of the input signal and algorithm parameters, such as input data window size, may need to be tuned to the specific signal of interest. Future designs of phase detection algorithms may benefit from explicitly calculating signal properties and using the information for adaptation (Van Zaen et al., 2010, Chen et al., 2013, Rodriguez Rivero and Ditterich, 2021).

## Author Contributions

ML and JYY conceived the project. ZML, SS, NZ and JYY performed the rodent neural recording experiments. ML performed simulation and analysis. ML and JYY wrote the manuscript.

## Acknowledgements

We thank Henry Jones, Department of Psychology, University of Chicago, for assisting with understanding the human EEG dataset.

## Supplemental data

**Figure S1.**
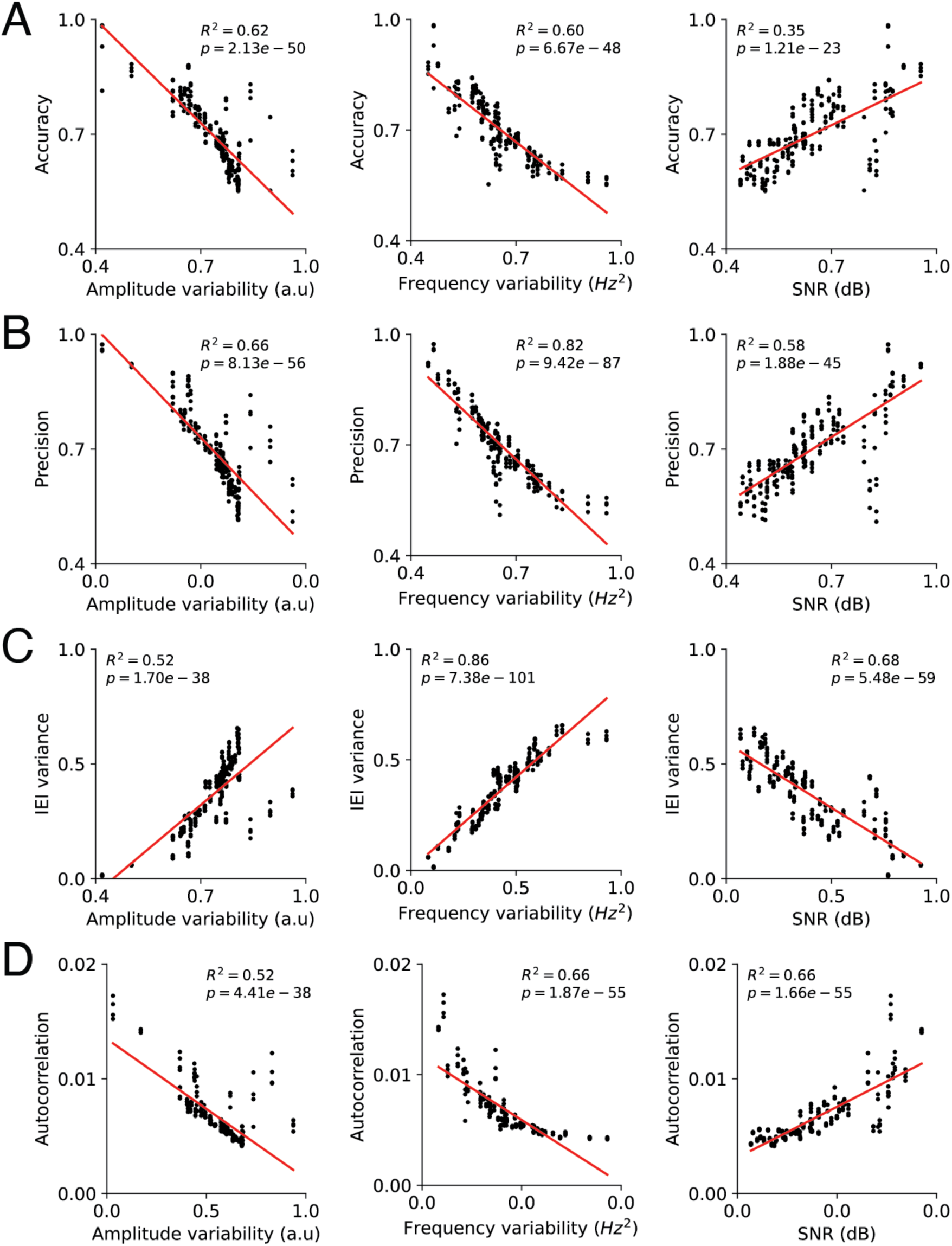
Linear regression between signal features and endpoint-corrected Hilbert transform performance metrics across all signal types. (A) Accuracy versus amplitude variability (left), frequency variability (center), and signal-to-noise ratio (right). (B) Precision versus amplitude variability (left), frequency variability (center), and signal-to-noise ratio (right). (C) IEI variance variance versus amplitude variability (left), frequency variability (center), and signal-to-noise ratio (right). (D) Autocorrelation versus amplitude variability (left), frequency variability (center), and signal-to-noise ratio (right).

**Figure S2.**
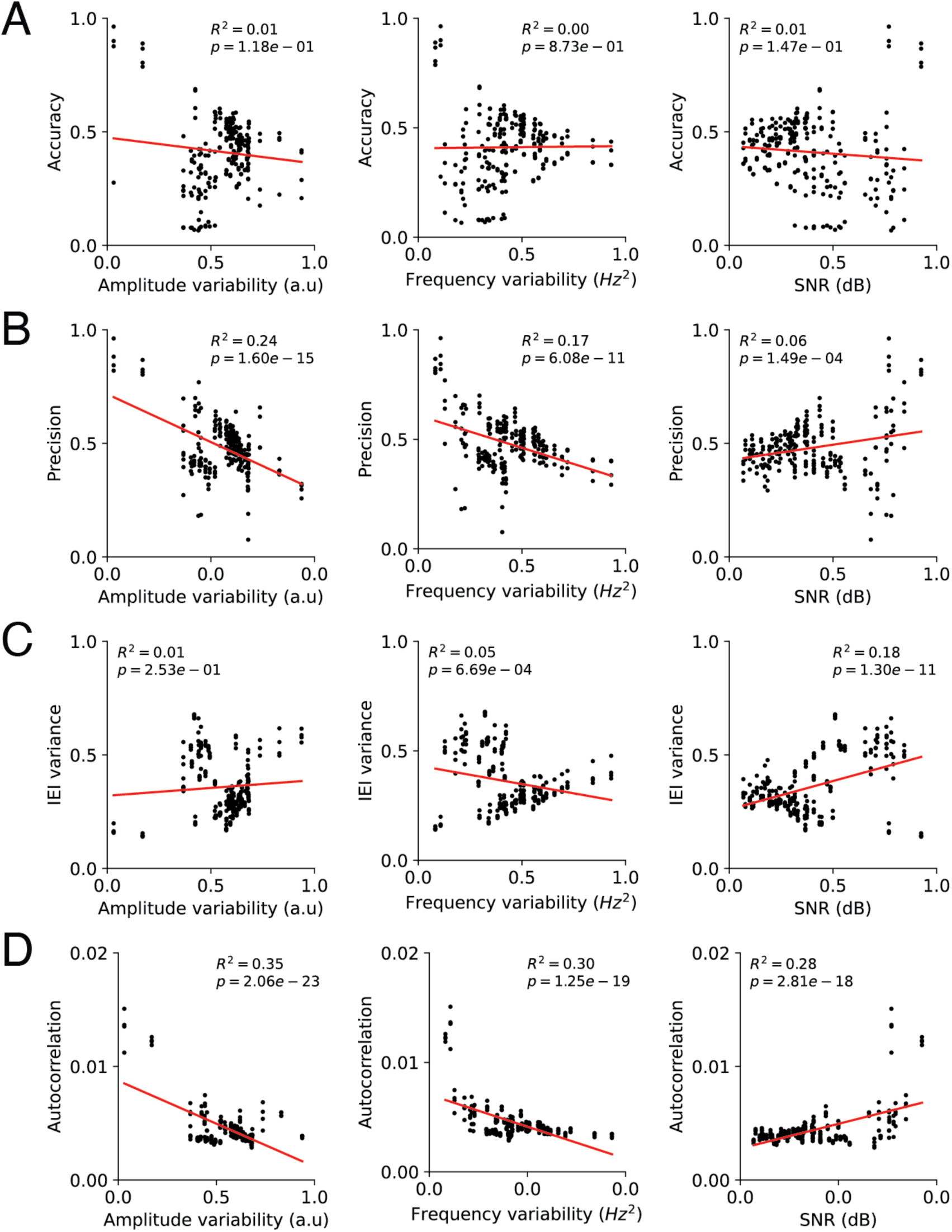
Linear regression between signal features and Hilbert transform algorithm performance metrics across all signal types. (A) Accuracy versus amplitude variability (left), frequency variability (center), and signal-to-noise ratio (right). (B) Precision versus amplitude variability (left), frequency variability (center), and signal-to-noise ratio (right). (C) IEI variance versus amplitude variability (left), frequency variability (center), and signal-to-noise ratio (right). (D) Autocorrelation versus amplitude variability (left), frequency variability (center), and signal-to-noise ratio (right).

**Figure S3.**
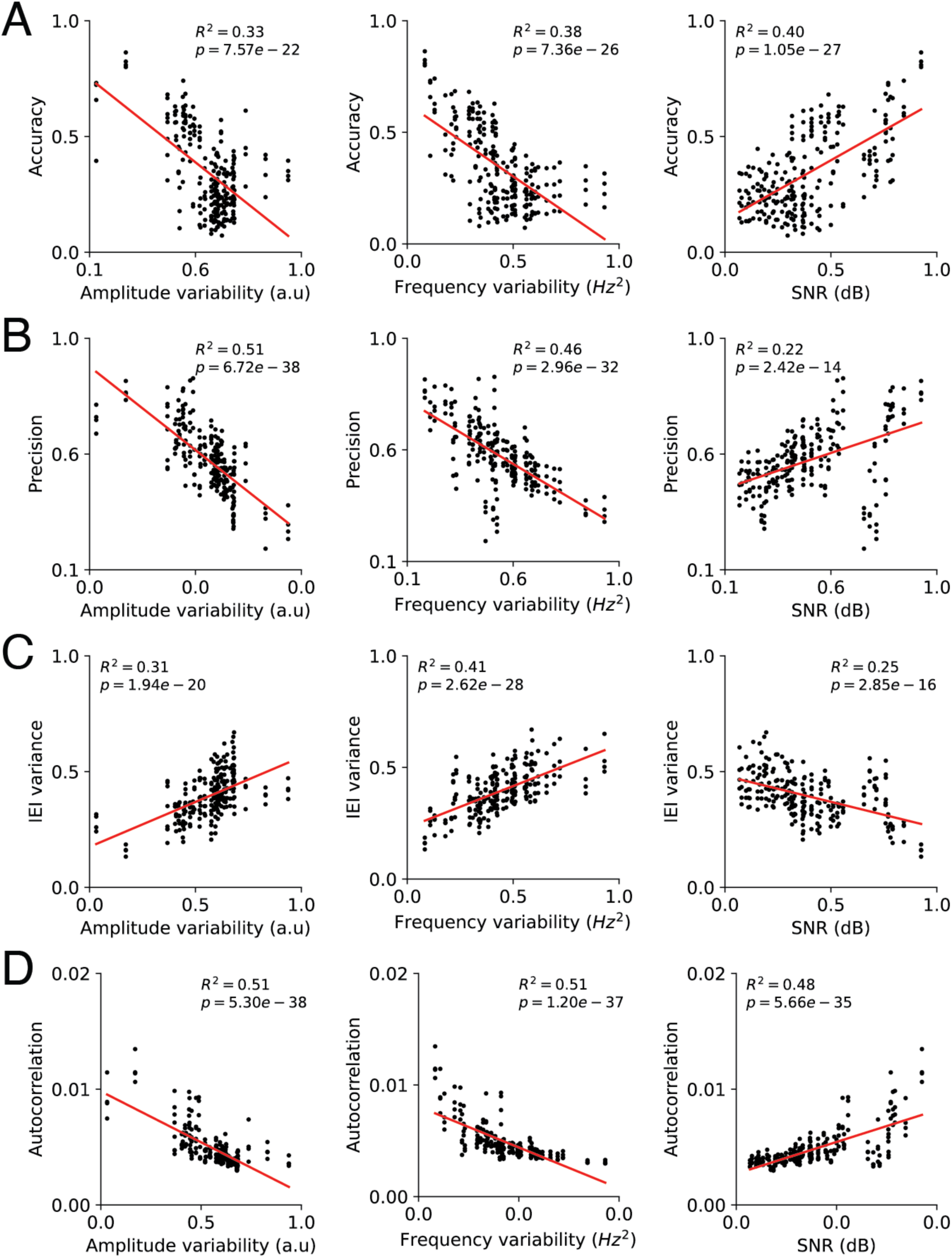
Linear regression between signal features and phase mapping algorithm performance metrics across all signal types. (A) Accuracy versus amplitude variability (left), frequency variability (center), and signal-to-noise ratio (right). (B) Precision versus amplitude variability (left), frequency variability (center), and signal-to-noise ratio (right). (C) IEI variance versus amplitude variability (left), frequency variability (center), and signal-to-noise ratio (right). (D) Autocorrelation versus amplitude variability (left), frequency variability (center), and signal-to-noise ratio (right).

**Table S1.** Two-way ANOVA of the effects of signal feature and algorithm on beta coefficients of the GLMs.

| *Source of variation* | *df* | *SS* | *MS* | *F* | *p-value* |
| --- | --- | --- | --- | --- | --- |
| *Signal features* | 2 | 0.8627 | 0.4314 | 6.0873 | 6.58e-03 |
| *Algorithm* | 2 | 0.3192 | 0.1596 | 2.2520 | 0.12 |
| *Interaction* | 4 | 0.2899 | 0.0725 | 1.0229 | 0.41 |

**Table S2.** Post-hoc comparison using Tukey’s Honest Significant Difference (HSD) test, pairwise comparison of beta coefficients between signal properties across algorithms and within each algorithm.

|  | p-value of Amplitude variability<br>vs. frequency variability | p-value of amplitude<br>variability vs. SNR | p-value of frequency<br>variability vs. SNR |
| --- | --- | --- | --- |
| Across algorithms | 0.281 | 5e-3 | 0.184 |
| ecHT | 9e-4 | 2.1e-6 | 6.5e-2 |
| HT | 0.721 | 0.858 | 0.272 |
| PM | 2.3e-3 | 0.917 | 0.913 |

**Table S3.** Two-way ANOVA of the effects of signal feature and algorithm on feature importance scores of the GLMs.

| *Source of variation* | *df* | *SS* | *MS* | *F* | *p-value* |
| --- | --- | --- | --- | --- | --- |
| *Signal features* | 2 | 3.1692 | 1.5846 | 14.3975 | 7.0e-6 |
| *Algorithm* | 2 | 1.7604 | 0.8802 | 7.9975 | 8.03e-04 |
| *Interaction* | 4 | 1.3851 | 0.3463 | 3.1462 | 2.01e-2 |

**Table S4.** Post-hoc comparison using Tukey’s Honest Significant Difference (HSD) test, pairwise comparison of feature importance scores between signal properties across algorithms and within each algorithm.

|  | p-value of Amplitude variability<br>vs. frequency variability | p-value of amplitude<br>variability vs. SNR | p-value of frequency<br>variability vs. SNR |
| --- | --- | --- | --- |
| Across algorithms | 0.445 | 1e-4 | 4.6e-3 |
| ecHT | 5.8e-6 | 4.23e-5 | 0.028 |
| HT | 0.952 | 6.7e-3 | 3.4e-3 |
| PM | 0.991 | 0.645 | 0.726 |

